# A brainstem-central amygdala circuit underlies defensive responses to learned threats

**DOI:** 10.1101/519249

**Authors:** Yiran Gu, Elena M. Vazey, Gary Aston-Jones, Longnian Lin, Joseph E. LeDoux, Robert M Sears

## Abstract

Norepinephrine (NE) plays a central role in the acquisition of aversive learning via actions in the lateral nucleus of the amygdala (LA)^1,2^. However, the function of NE in expression of aversively-conditioned responses has not been established. Given the role of the central nucleus of the amygdala (CeA) in the expression of such behaviors^3^, and the presence of NE projections in this brain nucleus, we assessed the effects of NE activity in the CeA on behavioral expression using receptor-specific pharmacology and cell-and projection-specific chemogenetic manipulations. We found that inhibition and activation of locus coeruleus (LC) neurons decreases and increases freezing to aversively conditioned cues, respectively. We then show that locally inhibiting or activating LC terminals in CeA is sufficient to achieve this bidirectional modulation of defensive reactions. These findings support the hypothesis that LC projections to CeA are required for the expression of defensive responses elicited by conditioned threats.

## Introduction

Much of the work describing the neural and behavioral mechanisms of defensive behavior and threat processing has used Pavlovian threat conditioning (PTC)^4^. This research has shown that the amygdala plays a crucial role in defensive reactions initiated by environmental threats^5,6^. PTC and the amygdala have both been implicated in fear and anxiety disorders^5,7^, as has the neuromodulator norepinephrine (NE). In the present study we explore the contribution of NE in the amygdala to the expression of amygdala controlled defensive behavior. During PTC, a neutral conditioned stimulus (CS; e.g., an acoustic tone) is paired with a noxious unconditioned stimulus (US; e.g., an electric foot shock) so that later presentation of the CS alone results in expression of defensive behaviors (the conditioned response; e.g., freezing). CS and US signals converge in both the lateral nucleus of the amygdala (LA) and the central nucleus of the amygdala (CeA)^5,8-10^. The LA communicates indirectly with CeA via the basal amygdala (BA)^9,11,12^, although a direct input to CeA has also been observed^11^. As a major output nucleus of the amygdala, the CeA coordinates defensive behavioral reactions and supports physiological adjustments in response to threatening stimuli via divergent projections to the midbrain^13-17^, lateral/paraventricular hypothalamus^15,18,19^, and medulla^3,13,20^.

NE is also implicated in fear and anxiety^21,22^. Aversive stimuli and stress increase levels of NE in the brain, including the amygdala^23^, largely through activation of the brain stem locus coeruleus (LC)^24,25^. LC stimulation or noxious stimuli (e.g., footshock) modulate the basolateral region of the amygdala (BLA)^23,24,26^, and studies suggest that this is through direct NE activity at β-adrenergic receptors (β-ARs)^2,27-30^. Notably, β-ARs in the LA are critical for initial acquisition (and indirectly for consolidation^2^ processes), but not expression of Pavlovian threat memories^1,2^, and in BLA (which includes LA and BA), for conditioned place aversion and anxiety-like behaviors^31^. Although much work on the role of NE in the amygdala has focused on the BLA, the CeA also receives significant NE inputs from the LC^32,33,34,35^. Despite this anatomical evidence, few studies have examined the contribution of NE inputs to CeA to the expression of defensive responses elicited by conditioned threats.

Here we describe the role of NE in the expression of Pavlovian threat memories and uncover key components of the underlying brain circuitry. We first show that CS-elicited defensive responses (freezing) decreased following systemic injection of the β-ARs antagonist (propranolol), while injection of β_2_-ARs agonist (procaterol) increased freezing. To test the role of LC in these adrenergic effects on behavioral expression, we used adeno-associated virus (AAV) vectors expressing DREADDs^36^ and engineered to target NE-expressing neurons in the LC (NE-LC)^19,37^. Inhibiting (hM4Di) or activating (hM3Dq) NE-LC neurons prior to the expression test by systemic Clozapine-N-oxide (CNO) injections caused reduced or enhanced behavioral expression, respectively. To test the hypothesis that CeA mediates this effect we used direct infusion of propranolol into CeA, which also reduced freezing. To test the role of a specific NE-LC➔CeA circuit in expression, we directly inhibited or activated axon terminals from the LC in CeA using intra-CeA CNO infusions prior to the expression test. Consistent with the pharmacology and NE-LC DREADDs studies, we found that inhibition and activation of LC terminals in CeA bidirectionally modulated freezing behavior. Finally, to test a requirement for β-ARs activation in the circuit-specific hM3Dq results, propranolol was co-infused with CNO in CeA. As predicted, propranolol attenuated CNO-induced enhancement of freezing. Taken together, these studies demonstrate that NE released from LC terminals in CeA enhances the expression of defensive responses elicited by learned threats.

## Methods

### Subjects

Adult male Sprague-Dawley rats were obtained from Hilltop Laboratory Animals, Inc. (Scottdale, PA, USA), weighing 250-275 g upon arrival. All animals were naive and allowed at least 1 week of acclimation to the vivarium before surgery and conditioning. Rats were individually housed in transparent plastic high-efficiency particulate absorption (HPEA)-filtered cages and maintained on a 12/12 h light/dark cycle (7:00 A.M.-7:00 P.M.) within a temperature-and humidity-controlled environment. Food and water were available *ad libitum* throughout the duration of the experiments. All experiments were conducted during the light cycle. All procedures were conducted in accordance with the *National Institutes of Health Guide for the Care and Use of Experimental Animals* and were approved by the New York University Animal Care and Use Committee.

### Stereotaxic Surgery

Rats were anesthetized with a mixture of ketamine (100 mg/kg, i.p.) and xylazine (10 mg/kg, i.p.), and placed in a stereotaxic apparatus (David Kopf Instruments, Tujunga, CA, USA). Supplemental doses of the mixture were given as needed to maintain a deep level of anesthesia. Brain areas were targeted using coordinates. After surgery, rats were administered buprenorphine hydrochloride (0.02 mg/kg, s.c.) for analgesia. And rats were given ketoprofen (5 mg/kg) for continuous 3 days to recover from surgery prior to behavioral manipulations.

For central amygdala (CeA) pharmacological infusion experiments, rats were bilateral implanted with stainless steel guide cannulae (22 gauge; Plastics One, Roanoke, VA, USA) were lowered into position aimed at the CeA (stereotaxic coordinates from bregma: anterior–posterior (AP) –2.8 mm, medial–lateral (ML) ±4.3 mm, dorsal–ventral (DV) –7.0 mm from skull) and secured to the skull using surgical screws and acrylic dental cement (Ortho-jet; Lang Dental Manufacturing Co.). Dummy cannulate (28 gauge) with extended 0.2 mm from the guides, were inserted to prevent clogging. During CeA pharmacology infusion, infusion cannulae (28 gauge) that extended 1.5 mm beyond the guides, were lowered into the CeA using the following coordinates from Paxinos and Watson.

For locus coeruelus (LC) viral injection experiments, DREADDs virus was bilaterally injected (stereotaxic coordinates from lambda: anterior–posterior (AP) –0.72 mm, medial–lateral (ML) ±1.35 mm, dorsal–ventral (DV) –7.5 mm from skull) to a volume of 1.4 µl/side using a 5.0 µl Hamilton Neuros syringe (Hamilton Co.) at a rate of 0.1 µl/min. After 3–6 weeks to allow for virus expression, animals were handled and subjected to behavioral conditioning as described below.

### Apparatus

For behavioral experiments, rats underwent threat conditioning in 1 of 6 identical chambers (Rat Test Cage; Coulbourn Instruments, Allentown, PA, USA) constructed of aluminum and Plexiglas walls, with metal stainless steel rod flooring that was attached to a shock generator (Model H13-15; Coulbourn Instruments). During habituation and threat conditioning, the chambers were lit with a single house light, and each chamber was enclosed within a sound isolation cubicle (Model H10-24A; Coulbourn Instruments; context A).

All the animals were tested in Med-Associate boxes without a house light. Testing took place in a modified context which consisted of dim red lighting, with smooth black plastic flooring, mild peppermint scent, mild lavender scent, and a striped pattern on the Plexiglas door (context B). An infrared digital camera, mounted on top of each chamber, allowed videotaping during behavioral procedures for offline subsequent behavioral scoring. A computer, installed with Graphic State 2 software and connected to the chambers via the Habitest Linc System (Coulbourn Instruments), delivered the presentation of tone stimuli during behavioral sessions.

### Viral vectors

Excitatory (hM3Dq: AAV9/PRS×8-HA-hM3Dq-SV40-PolyA), inhibitory (hM4Di: AAV9/PRS×8-HA-hM4Di-SV40-PolyA) and control vectors (Control: AAV9/PRS×8-mCherry-WPRE-rBG) were subcloned by Dr. Elena M. Vazey from Gary Aston-Jones’ lab and packaged by the University of Pennsylvania Vector Core. The synthetic PRS×8 promoter was used to restrict expression of the hM3Dq/hM4Di DREADDs to noradrenergic neurons in the locus coeruleus (LC).

### Drug Preparation and Microinfusion

Propranolol ((±)-Propranolol hydrochloride) and Procaterol (Procaterol hydrochloride) were obtained from Sigma-Aldrich Co. (St. Louis, MO, USA), and freshly dissolved in 0.9% sterile saline immediately prior to injections. For intraperitoneal injection experiments, concentrations for propranolol and procaterol were 10 mg/kg and 300 µg/kg, respectively. For CeA microinfusion experiments, propranolol was dissolved in 0.9% sterile saline and administered at 1.0 µg /0.3 µl. CNO for IP experiments was obtained from the NIH as part of the Rapid Access to Investigative Drug Program funded by the NINDS and prepared in a 7% DMSO+ 0.9% sterile saline, which was also used for vehicle infusion. For intracranial infusions, CNO from Sigma-Alrdich was dissolved in 0.9% sterile saline.

For intracranial drug infusions, internal infusion cannulae were attached to 10 µl Hamilton syringes via 0.015” × 0.043” × 0.014” polyethylene tubing obtained from A-M Systems, Inc. (Carlsborg, WA, USA). Tubing and syringes were backfilled with distilled water, and a small air bubble was introduced to separate the water from the infusate. Rats were bilaterally infused with 0.3 µl using an infusion pump (PHD 2000; Harvard Apparatus) at a constant rate of 0.1 µl/min. Animals were allowed to move freely in their home cage during infusions. After infusion was complete, cannulae were left in place for an additional 1-2 mins to allow drug diffusion away from the cannula tip.

### Pavlovian threat conditioning and testing procedures

All rats were habituated in groups of six rats at a time on Day 1, during which rats were free to explore the conditioning box (context A) for thirty minutes. After habituation, rats were returned to their home cages within the colony room.

Conditioning was conducted in groups of six rats at a time, each in a different chamber (context A). Following an initial 5-min acclimation period, all rats were presented with three conditioning trials (CS–US pairings) on Day 2. The CS was a 30-s, 5 kHz, 80 dB SPL sine-wave tone, which co-terminated with a 1-s, 0.6 mA standard footshock US, or a 0.4 mA weak footshock US delivered through the rod flooring. The mean inter-trial interval was 4 min (2–6 min range) for both conditioning and testing sessions. After conditioning, rats were returned to their home cages within the colony room.

Expression testing for CS-elicited threat was conducted in the modified context (context B) on Day 3, 1 day after conditioning. After the 5-min acclimation period, rats were presented with five CS-alone presentations, using the same stimulus parameters as in conditioning, but excluding the footshock US. Behavior was recorded and freezing was scored as described below.

Drug-free testing for CS-elicited freezing was conducted in the modified context (context C) on Day 4, 2 days after conditioning. After the 5-min acclimation period, rats were presented with five CS-alone presentations, using the same stimulus parameters as in conditioning, but excluding the footshock US. Behavior was recorded and freezing was scored as described below.

### Measurement of Freezing Behavior

Freezing was used to measure the conditioned threat response and was defined as the cessation of all movement with the exception of respiration-related movement and non-awake or rest body posture^38^. Behavior was videotaped and later scored offline with a digital stopwatch by recording the total time spent freezing during every 30-s tone CS. Pre-CS freezing was also scored during the 30-s interval prior to the initial tone onset, and was used as a measure of non-specific freezing to the context. Freezing was scored by two experimenters blind to drug group allocation and the data averaged.

### Histology and immunohistochemistry

Following behavior experiments, animals were overdosed with 25% chloral hydrate and transcardially perfused with either 10% formalin for histology to assess cannula placement or 4% paraformaldehyde in 0.01 M PBS for immunohistochemistry.

Tissue processed for cannula placement was post-fixed in 10% formalin at 4 °C until prepared for histological staining. For immunohistochemical processing, tissue was cryoprotected in a 30% sucrose– 4% paraformaldehyde solution (in 0.01 M PBS) at 4 °C for at least 1 day and then stored in 0.01 M PBS. Brains were blocked coronally and cut on a freezing microtome.

For histological verification of cannula targeting, tissue was cut at a thickness of 50 μm and kept in 0.01 M PBS until mounted on gelatin-coated slides and dried overnight. After standard histological Nissl staining and coverslipping, sections were examined on a light microscope for injector tip localization into the central amygdala. Only data from rats with bilateral injector placements localized to the central amygdala were included in the study.

For immunohistochemistry, tissue was cut at 35 μm. Before antibody incubation, floating tissue was rinsed with agitation three times in 0.01 M PBS and blocked in 1% BSA in 0.01 M PBS for 1 h at room temperature. Immunohistochemical detection was achieved in primary antibody solutions containing 1% BSA, 0.4% Triton-X 100, and 0.02% NaAz.

DREADDs animal brain sections were incubated overnight at room temperature in rabbit anti-HA (for detection of HA-Tag; 1:500; Cell Signaling Technology, MA) and mouse anti–dopamine beta hydroxylase (DBH) (1:2000; EMD MIllipore, MA) antibodies for verification of viral expression in LC neurons, long-projection terminals in CeA and cell specificity of viral expression. Virus control animal brain sections were incubated overnight at room temperature in rabbit anti-DsRed (for detection of mCherry; 1:500; Clontech Laboratories, CA) and anti–dopamine beta hydroxylase (DBH) (1:2000; EMD MIllipore, MA) antibodies.

Following primary antibody incubation, sections were rinsed with agitation three times for 5 min in 0.01 M PBS and incubated in Alexa Fluor goat anti-rabbit 594 or goat anti-mouse 488 secondary antibody (1:200; Life Technologies, CA) in 0.01 M PBS. Sections were rinsed three times for 5 min in PBS, mounted on gelatin-coated slides, and allowed to dry for several hours, followed by a brief wash in ddH_2_O to remove excess salt (PBS), coverslipped in aqueous mount (ProLong Gold Antifade Reagent with DAPI; Life Technologies, CA), and allowed to cure overnight at room temperature before fluorescence imaging. Leica TCS SP8 confocal microscope (Leica) or Olympus VS120 fluorescent microscope (Olympus) were used to capture images. Imaging data were processed and analyzed with ImageJ software (NIH). For all experiments, animals were excluded from analysis if virus-expression was insufficient or cannulae targeting was outside the areas of interest.

### Statistics

For behavioral data with two groups, a Student’s t test was used to analyze freezing levels (baseline or CS-elicited freezing). One-way ANOVA was used for comparing more than one group followed by a Tukey’s multiple comparison tests. Mean CS freezing data between drug-treatment days and drug-free days was analyzed using repeated measures ANOVAs followed by Sidak multiple comparisons tests. Bartlett’s test for equal variances was used for one-way ANOVAs, whereas the F test was used for the Student t test to confirm that variances were not significantly different in compared groups. Error bars in all figures represent ± SEM. Data were analyzed using GraphPad Prism.

## Results

We first investigated the contribution of β-ARs activity to the expression of Pavlovian conditioned defensive responses. Twenty-four hours following cued threat conditioning using a moderate protocol (0.6 mA shock), rats were administered the β-ARs antagonist propranolol (10 mg/kg, n = 12) or saline (n = 12) and tested for freezing responses to the CSs (Figure 1a). Propranolol significantly attenuated CS-evoked freezing levels (t (22) = 3.894, ***p = 0.0008), and slightly reduced baseline freezing (t (22) = 2.348, *p = 0.0283) (Figure 1b, left panel), effects that were not observed during a drug-free test in the same animals (Figure 1b, center panel). In separate groups of animals, propranolol still reduced freezing with stronger training (Supplementary Figure 1a, 1b, n=11/group, 1.0 mA shock training (t (20) = 2.129, *p = 0.0459)) and during a remote memory test (Supplementary Figure 1c, 1d, n=9/group, 1 month after conditioning (BL freezing: Student’s t test, t (16) = 2.297, *p = 0.0354, CS freezing: t (16) = 5.930, ****p < 0.0001).

**Figure 1.**
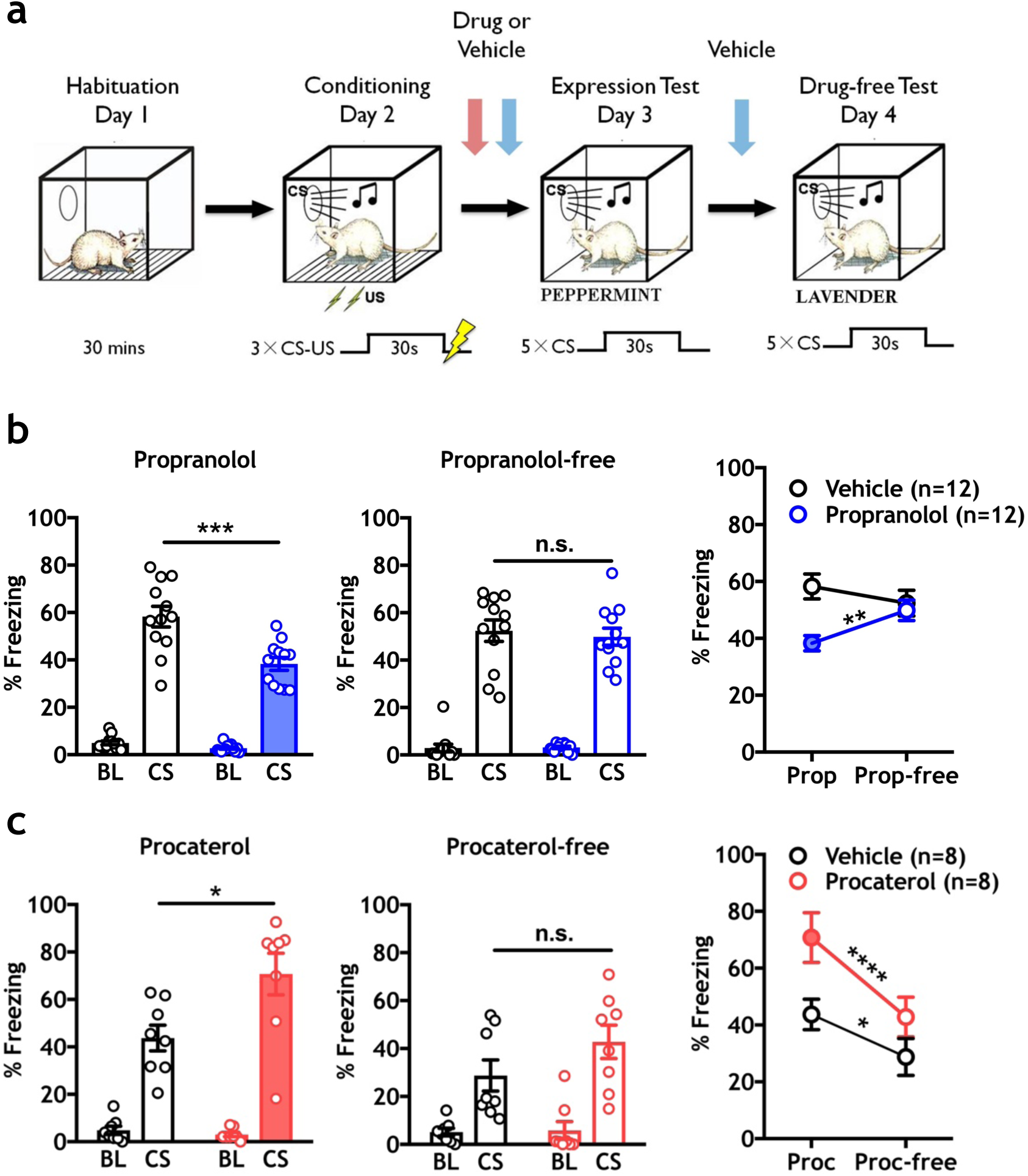
Norepinephrine β-ARs activity is required for CS-elicited freezing in threat-conditioned animals. **a.** Experimental timeline depicting habituation, training (0.6 mA US), expression test (Day 3) and drug-free expression test (Day 4) phases. Vertical arrows indicate time of drug (red arrow) or vehicle (blue arrow) injection for each manipulation. **b.** Propranolol (10 mg/kg) reduced baseline (*p = 0.0283) and CS-elicited (***p = 0.0008) freezing levels during the expression test compared to vehicle control animals (left panel), with no effect observed between groups during a drug-free test (center panel). A within-subject comparison of propranolol treatment versus propranolol-free treatment on CS-elicited freezing showed a significant difference for drug treatment (two-way RM ANOVA test, Interaction: F (1, 22) = 15.79, ***p = 0.0006; Time (Drug vs. Drug-free) F (1,22) = 1.678, p = 0.2086; Drug vs. Vehicle F (1, 22) = 5.021, *p = 0.0355; Sidak MCS, **p < 0.01 between days for propranolol treated animals, n.s. for vehicle-treated animals). **c.** Injection of the specific β_2_-ARs agonist procaterol (300 µg/kg) enhanced CS-elicited freezing during the expression test (n = 8/group; left panel, *p = 0.0200), with no effect during a drug free test (center panel). Within subject analysis showed a main effect of procaterol on CS-elicited freezing between days in both groups (two-way RM ANOVA test, Interaction: F (1, 14) = 3.991, p = 0.0655; Drug: F (1, 14) =4.786, *p = 0.0462; Time (Drug vs. Drug-free), F (1, 14) = 43.73, ****p < 0.0001; Sidak MCS, ****p < 0.0001 between days for procaterol treated animals, *p < 0.05 for vehicle-treated animals). All error bars indicate mean ± SEM. *p < 0.05, **p< 0.01, ***p< 0.001, ****p < 0.0001.

To assess the effects of β_2_-receptor activation on memory expression, two groups of rats were administered systemic injections of the β_2_-receptor agonist procaterol (300 µg/kg, n = 8) or saline vehicle (n = 8) prior to the expression test. Procaterol significantly increased CS-evoked freezing levels (Figure 1c left panel, 300 µg/kg, (t (14) = 2.625, *p = 0.0200)), with no difference observed between groups when the drug was not onboard (Figure 1c, center panel). Collectively, these data reveal that β-ARs positively modulate the expression of defensive responses regardless of memory strength or time since memory formation, and blockade or activation does not have long-term effects on behavioral plasticity.

### Chemogenetic inhibition of LC attenuates freezing to a conditioned cue

LC neurons send NE efferent throughout the brain, including the amygdala^32-35^. We therefore tested whether LC activity is necessary for the expression of defensive responses. Using AAV vectors expressing DREADDs (hM4Di) or a fluorescent reporter (mCherry) under the control of a synthetic promoter (PRS × 8)^19,37,39^, we observed expression restricted to NE (dopamine β hydroxylase (DBH)-positive)) neurons in LC (Figure 2b). Following bilateral AAV injection in LC, an hM4Di group (n = 9) and an mCherry (n = 7) group were trained using a moderate protocol (3 CS-US pairings, 0.6 mA US), and a third hM4Di group (n=7) received 3 CS presentations without footshock to control for non-specific effects on freezing behavior. All rats received systemic injections of CNO (5.0 mg/kg) prior to the expression test (Figure 2a). CNO significantly attenuated CS-elicited freezing levels only in conditioned animals expressing hM4Di compared to conditioned animals expressing mCherry alone (CS freezing: One-way ANOVA F (2, 20) = 126.4, ****p < 0.0001, Tukey’s MCT: hM4Di untrained vs. mCherry trained, ****p < 0.0001, hM4Di untrained vs. hM4Di trained, ****p < 0.0001, mCherry trained vs. hM4Di trained, **p < 0.01), with no effect observed in the untrained behavioral controls expressing hM4Di (Figure 2c, left panel). No significant effect was observed between trained groups in freezing levels during a CNO-free test, and freezing remained negligible in the untrained group (Figure 2c, center panel). These data suggest that LC-NE activity positively modulates the expression of learned threat reactions.

**Figure 2.**
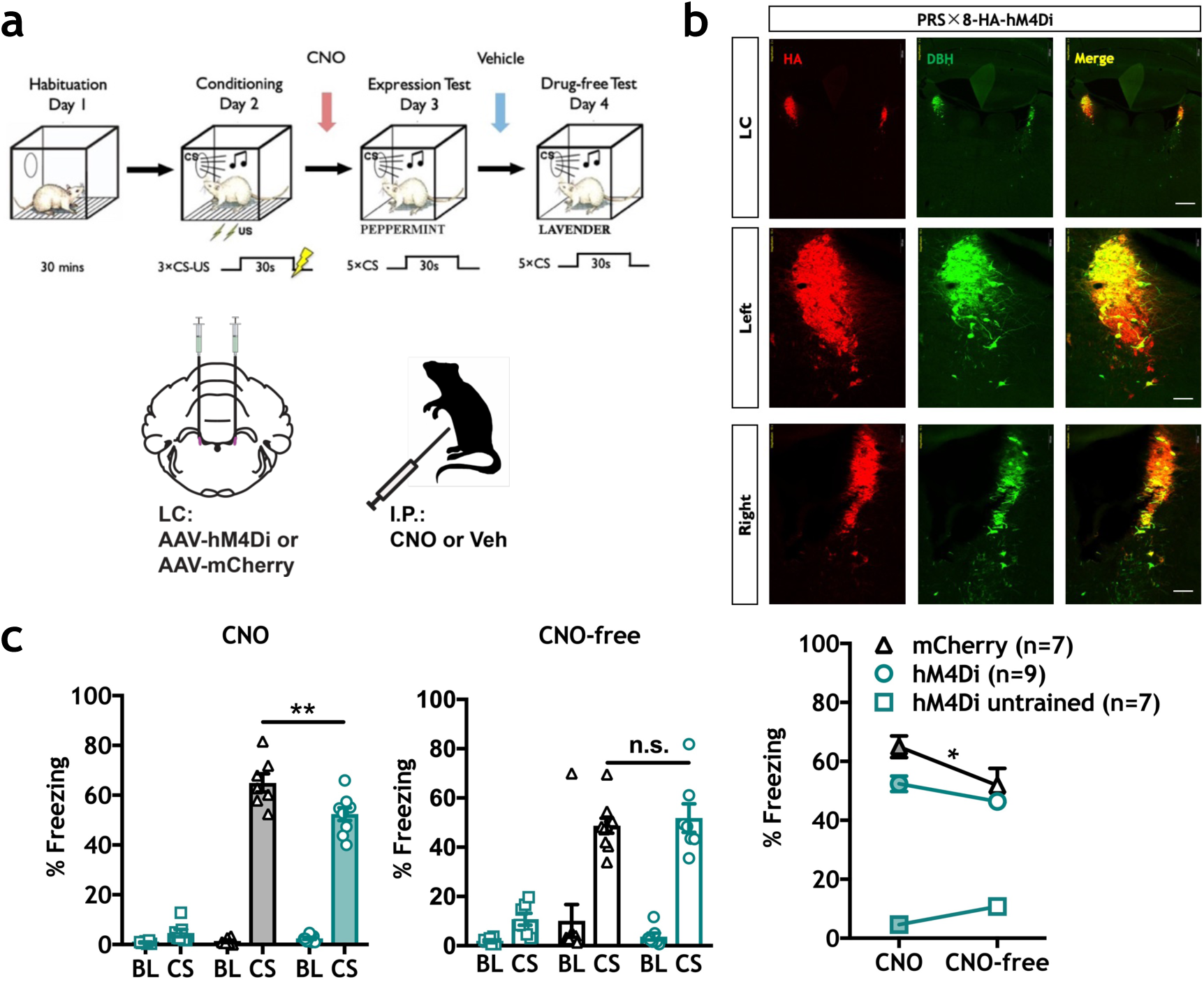
Chemogenetic inhibition of LC-NE signaling decreases CS-elicited freezing. **a** Top: Timeline indicating habituation, conditioning (0.6 mA US), expression test (Day 3) and drug-free test (Day 4) phases. Bottom: Schematic depicting hM4Di or mCherry virus injection and CNO treatment strategy. **b.** Representative immunohistochemistry (IHC) images show robust and selective targeting of hemagglutinin-tagged (HA) hM4Di receptors to DBH^+^ LC neurons. (Red = HA; Green = dopamine β hydroxylase (DBH); Yellow = indicates co-localization. Scale bars: top three panels = 500 µm, middle and bottom six panels = 100 µm. **c.** On conditioning day, hM4Di paired (n=9) and mCherry paired groups (n=7) were threat conditioned, and an unpaired hM4Di control group (n=7) received three tones alone. CNO (5.0 mg/kg) inhibition of LC-NE neurons significantly decreased CS-elicited freezing in trained hM4Di animals compared to mCherry controls (one-way ANOVA, F (2,20) = 126.4, ****P < 0.0001; Tukey’s MCS, **P < 0.01), with no difference observed between hM4Di paired and mCherry paired groups during the drug-free test. Within subject analysis revealed a slight reduction in CS-elicited freezing between days in the mCherry group, with no significant reduction in the hM4Di paired group (two-way RM ANOVA test, Interaction: F (2, 20) = 4.236, *p = 0.0292; Training X virus: F (2, 20) = 134.5, ****p < 0.0001; Time (CNO vs. CNO-free): F (1,20) = 2.692, p = 0.1165; Sidak MCS, CNO vs. CNO-free: hM4Di untrained, p = n.s., hM4Di trained, p = n.s., mCherry *p < 0.05). All error bars indicate mean ± SEM. *p< 0.05, **p< 0.01, ****p < 0.0001.

### Chemogenetic activation of LC enhances freezing to a conditioned cue

Next, excitatory DREADDs (hM3Dq) were used to determine how stimulation of LC-NE activity would affect CS-elicited freezing. hM3Dq-and mCherry-immunopositive neurons were observed to co-localize with the DBH-positive neurons in LC^19,37^, red fluorescence was detected throughout the entire locus coeruleus but not in neighboring noradrenergic or dopaminergic regions (Figure 3b, Supplementary Figure 4), which showed that viral targeting of the locus coeruleus was specific. An hM3Dq group (n = 9) and an mCherry group (n = 7) were trained using a weak protocol (to avoid ceiling effects, 3 CS-US pairings, 0.4 mA shock), and a third hM3Dq group (n = 7) received 3 CS-alone presentations without footshock to control for unconditioned freezing behaviors. Systemic CNO (1.0 mg/kg) significantly increased CS-elicited freezing levels in trained hM3Dq-expressing animals compared with mCherry-expressing animals (F (2, 20) = 69.54, ****p < 0.0001, Tukey’s MCT: hM3Dq untrained vs. mCherry trained, ****p < 0.0001, hM3Dq untrained vs. hM3Dq trained, ****p <0.0001, mCherry trained vs. hM3Dq trained, ***p < 0.001), with minimal CS-elicited freezing responses observed in the untrained control group (Figure 3c, left panel). No significant effects were observed between trained groups during a CNO-free test (Figure 3c, center panel). Taken together, these and hM4Di data (Figure 2) suggest that LC activity positively modulates CS-elicited freezing.

**Figure 3.**
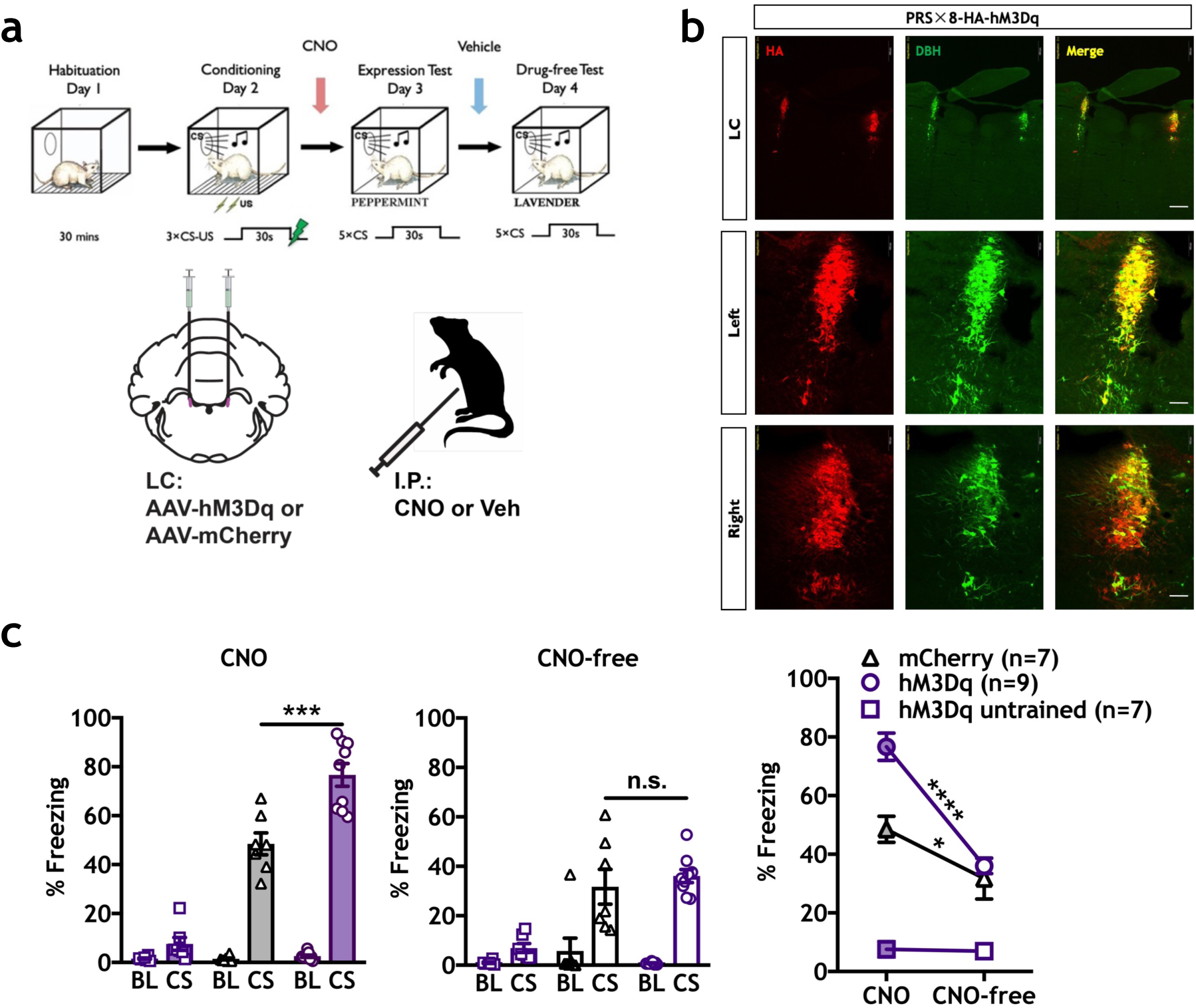
Chemogenetic activation of LC-NE signaling increases CS-elicited freezing. **a.** (Top) Timeline indicating habituation, mild conditioning (0.4 mA US), expression test and drug-free test phases. (Bottom) Schematic depicting hM3Dq or mCherry virus injection and CNO treatment strategy. **b.** Representative IHC images show robust and selective targeting of hM3Dq-HA to DBH^+^ LC neurons. (Red = HA; Green = DBH; Yellow = co-localization). Scale bars: top three panels = 500 µm, middle and bottom six panels = 100 µm. **c.** Systemic injection of CNO (1.0 mg/kg) prior to the expression test significantly enhanced freezing in the trained hM3Dq group (n=9) compared to the trained mCherry group (n=7; left panel, one-way ANOVA, F (2,20) = 69.54, ****P < 0.0001, Tukey’s MCS, ***P<0.001). No differences were observed between groups during a CNO-free expression test (center panel). A difference in CS-elicited freezing was observed in trained animals between CNO-and CNO-free tests (two-way RM ANOVA test, Interaction: F (2, 20) = 16.62, ****p < 0.0001; Training X virus: F (2, 20) = 55.64, ****p < 0.0001, Time (CNO vs. CNO-free): F (1,20) = 43.45, ****p < 0.0001; Sidak MCS, CNO vs. CNO-free: hM4Di untrained, p = n.s., hM3Dq trained, ****p < 0.0001, mCherry trained, *p < 0.05). All error bars indicate mean ± SEM. *p< 0.05, ***p< 0.001, ****p < 0.0001.

### Pharmacological blockade of β-ARs in CeA attenuates freezing

We found that pharmacological and chemogenetic manipulation of brain NE activity bidirectionally modulated behavioral expression. We next test the hypothesis that NE activity in the CeA positively modulates CS-elicited freezing. Rats received bilateral microinjections of the β-ARs antagonist propranolol or vehicle in the CeA prior to the expression test. Results show that propranolol (1.0 µg/0.3µL/side) significantly attenuated CS-elicited freezing levels compared to vehicle controls (n = 7/group, t (12) =5.324, ***p = 0.0002) (Figure 4a, right panel). Therefore, β-ARs activity specifically in CeA mediates the expression of responses to learned threats.

**Figure 4.**
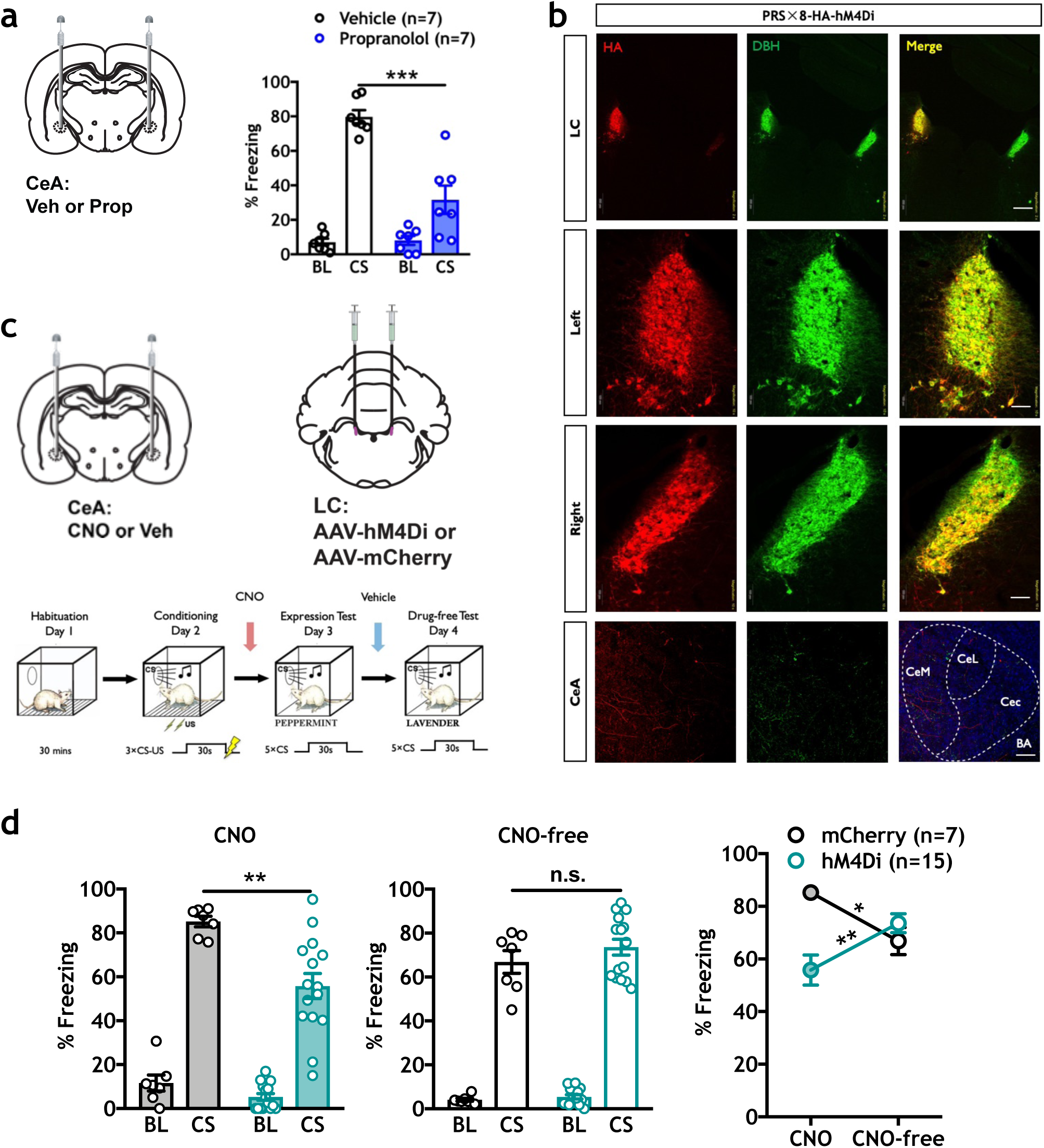
CeA blockade of β-ARs or chemogenetic inhibition of LC-NE axon terminals in the CeA decreases CS-elicited freezing. **a.** Propranolol infusions in CeA (1.0 µg/0.3 µl/side) significantly reduced CS-elicited freezing (***p = 0.0002). **b.** Robust and selective targeting of hM4Di-HA to DBH^+^ LC neurons and detectable expression in axons projecting to CeA after six weeks (Red = HA; Green = DBH; Yellow = co-localization; Blue = DAPI). Scale bars: top three LC panels = 500 µm, middle six LC panels = 100 µm, bottom three CeA panels = 50 µm. **c.** Virus and CNO infusion strategy and experimental timeline. **d**. CNO infusions in CeA (1.0 µM/0.3 µl/side) significantly reduced CS-elicited freezing in hM4Di animals (n=15) compared to the mCherry (n=7) group (t (20) =3.401, **p = 0.0028). No differences were observed between groups during the CNO-free expression test (center panel). Differences were observed in CS-elicited freezing in both groups between CNO-and CNO-free tests (two-way RM ANOVA test, Interaction: F (1, 20) = 20.13, ***p = 0.0002; Virus: F (1, 20) = 3.088, p = 0.0942; Time (CNO vs. CNO-free): F (1,20) = 0.005266, p = 0.9429; Sidak MCS, CNO vs. CNO-free: mCherry, *p < 0.05, hM4Di, **p < 0.01). All error bars indicate mean ± SEM. *p< 0.05, **p< 0.01, ***p< 0.001. Central medial amygdala (CeM), Central lateral amygdala (CeL), Central capsular amygdala (Cec), Basal amygdala (BA).

### Chemogenetic blockade of LC terminals in CeA attenuates freezing

We have shown that NE neurons in LC and β-ARs activity in CeA play a critical role in defensive responses. To test the LC➔CeA circuit, we combined targeted expression of hM4Di in LC-NE with direct CNO infusions to block axon terminal activity in CeA (Figure 4b). Six weeks following surgery, animals were trained using a moderate conditioning protocol (Figure 4c) and memory tested the subsequent day. Pre-test infusion of CNO in CeA (1.0 µM/0.3 µl/side) in hM4Di-expressing animals significantly attenuated freezing levels compared to mCherry controls (Figure 4d left panel, n=7-15/group, t (20) =3.401, **p = 0.0028). During a CNO-free test, intra-CeA infusion of vehicle did not significantly affect freezing behavior between groups (Figure 4d, center panel). These data support the hypothesis that LC➔CeA circuit activity is necessary for the expression of conditioned threat reactions.

### Chemogenetic activation of LC-NE terminals in CeA enhances CS-elicited freezing

To further confirm a role for a LC➔CeA circuit in defensive responses, we combined targeted expression of hM3Dq in LC-NE with direct CNO infusions to activate axon terminals in CeA. Six weeks following surgery, animals were trained using a mild conditioning protocol followed by an expression test the next day (Figure 5c). Following bilateral microinjections of CNO in CeA (1.0 µM/0.3 µl/side), we found that CNO significantly increased freezing levels in hM3Dq animals compared with the mCherry control group (t (16) =3.114, **p = 0.0067). (Figure 5c, left panel, n=8-10/group). In a subsequent CNO-free test, CS-elicited freezing was not different between groups (Figure 5c, center panel).

**Figure 5.**
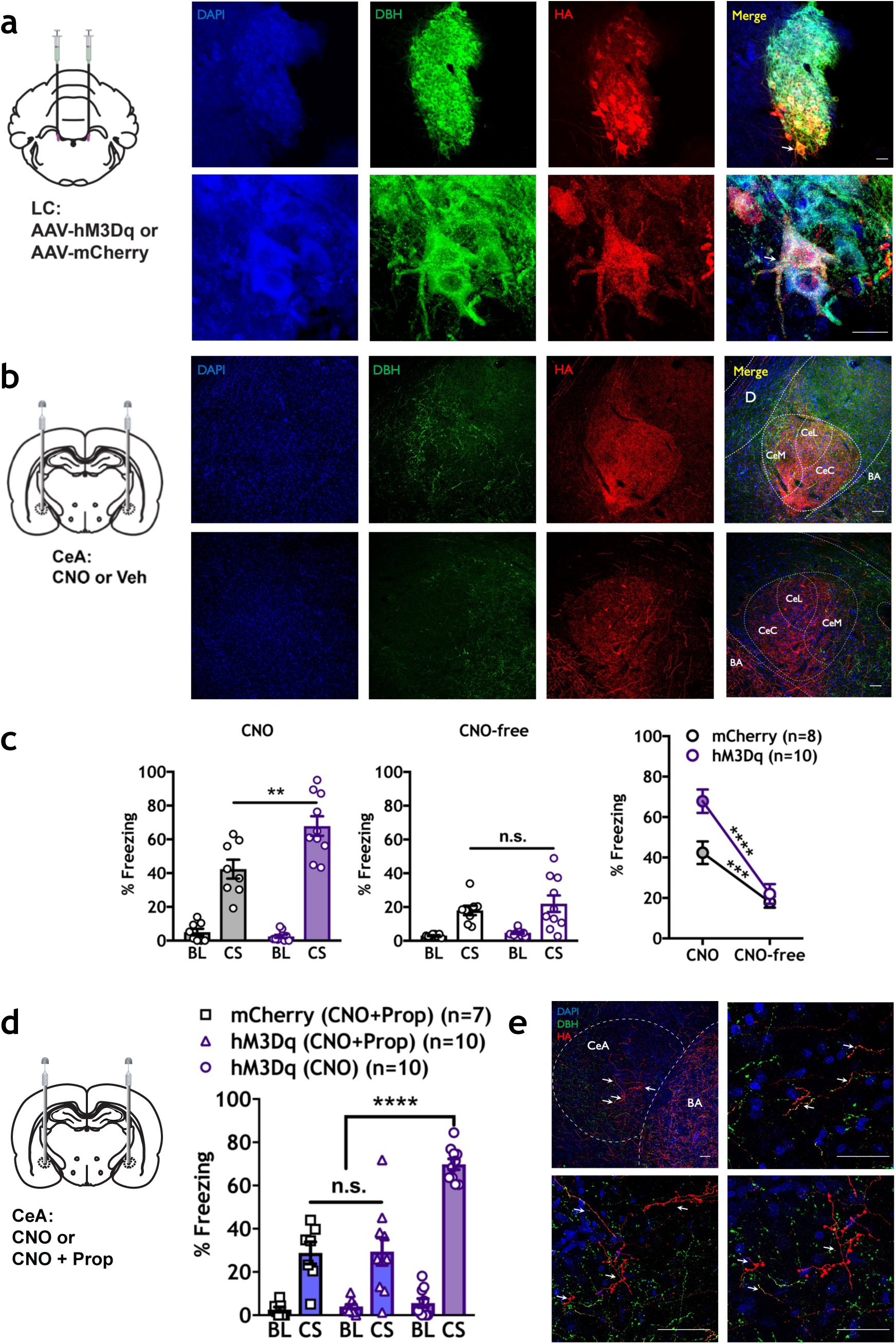
Chemogenetic activation of LC-NE terminals in CeA increases CS-elicited freezing behavior and is blocked by a β-AR antagonist. **a.** hM3Dq (HA) expression in LC (Red = HA; Green = DBH; Yellow = co-localization; Blue = DAPI). Arrow indicates the magnification of the same LC neuron. Scale bars: LC panels = 50 µm, respectively. **b.** HA-immunopositive terminals from LC were detected in CeA. Scale bars: CeA panels = 50 µm. **c.** Infusions of CNO (1.0 µM/0.3 µl/side) led to a significant increase in CS-elicited freezing in hM3Dq (n = 10) animals compared to mCherry controls (left panel, n = 8, t (16) =3.114, **p = 0.0067), with no difference observed between groups during a CNO-free test (center panel). Differences were observed in CS-elicited freezing in both groups between CNO-and CNO-free tests (two-way RM ANOVA test, Interaction: F (1, 16) = 11.78, **p = 0.0034; Virus: F (1, 16) = 5.197, *p = 0.0367; Time (CNO vs. CNO-free): F (1,16) = 125.7, ****p < 0.0001; Sidak MCS, CNO vs. CNO-free: mCherry, ***p < 0.001, hM3Dq, ****p < 0.0001). **d.** Prior to the expression test, animals received intra-CeA infusions of CNO alone (hM3Dq (CNO); (n=10)), or a cocktail of CNO and propranolol (1.0 µM/0.3 µl/side + 1.0 µg/0.3 µl/side) in hM3Dq (n=10) and mCherry (n=7) animals. Propranolol significantly reduced the effect of CNO in hM3Dq-expressing, propranolol-treated animals (one-way ANOVA, F (2, 24) = 23.77, ****P<0.0001, Tukey’s MCS, mCherry (CNO+Prop) vs. hM3Dq (CNO), ****p < 0.0001 and hM3Dq (CNO+Prop) vs. hM3Dq (CNO), ****p < 0.0001). **e.** Representative IHC images show robust and selective targeting of hM3Dq-HA to DBH^+^ LC neurons and strong expression in CeA terminals. (Red = HA; Green = DBH; Yellow = overlapping indicates example co-localization; Blue = DAPI.) Scale bars, CeA panels = 50 µm, respectively. Arrow indicates the magnification of the LC-NE terminals in CeA. All error bars indicate mean ± SEM. **p < 0.01, ****p < 0.0001. Central medial amygdala (CeM), Central lateral amygdala (CeL), Central capsular amygdala (Cec), Basal amygdala (BA).

### Enhancement of CS-elicited freezing by chemogenetic activation of the LC➔CeA circuit requires β-ARs activity in CeA

Next, we determined if chemogenetic enhancement of CS-elicited freezing requires NE activity at β-ARs in CeA. Prior to the expression test, hM3Dq rats were administered bilateral microinjections of either CNO alone (1.0 µM/0.3 µl/side) or a cocktail of propranolol (1.0 µg/0.3 µl/side) and CNO (1.0 µM/0.3 µl/side) in CeA. An mCherry group was also administered the propranolol-CNO cocktail. Results showed that CNO alone significantly enhanced CS-elicited freezing in hM3Dq animals, whereas inhibition of β-ARs in CeA significantly reduced this effect (one-way ANOVA: F (2, 24) = 23.77, ****p < 0.0001, Tukey’s MCS: mCherry (CNO+Prop) vs. hM3Dq (CNO+Prop), n.s., mCherry (CNO+Prop) vs. hM3Dq (CNO), ****p < 0.0001, hM3Dq (CNO+Prop) vs. hM3Dq (CNO), ****p < 0.0001) (Figure 5d, n=7-10/group). No difference was observed between hM3Dq (CNO + Prop) and mCherry (CNO + Prop) groups. These data suggest that freezing reactions require NE release from LC terminals in CeA.

## Discussion

Here we establish neural underpinnings of noradrenergic modulation of Pavlovian defensive reactions. First, we found that propranolol-mediated blockade of β-ARs significantly reduced behavioral expression of PTC (Figure 1b, S1b, S1d), whereas systemic activation of β_2_-ARs using procaterol enhanced it (Figure 1c). Endogenous noradrenergic activity in behavioral expression was then confirmed through chemogenetic reduction (Figure 2c) or enhancement (Figure 3c) of noradrenergic LC activity. Furthermore, blockade of β-ARs in the CeA, the major output nucleus of the amygdala mediating defensive reactions (Figure 4a) or inhibition of the LC➔CeA circuit (Figure 4d) yielded results consistent with systemic propranolol (Figure 1b) and chemogenetic LC inhibition (Figure 2c). As with procaterol injections (Figure 1c) and LC activation (Figure 3c), stimulation of LC➔CeA inputs enhanced CS-elicited freezing (Figure 5c). This enhancement was blocked by direct infusion of propranolol, suggesting that NE released from LC axons acts through β-ARs in CeA (Figure 5d). Taken together, these data suggest that LC➔CeA projections release NE into the CeA where β-ARs activation positively modulates the degree of CS-elicited freezing.

Noradrenergic LC neurons phasically respond to salient stimuli (learned or novel) in all sensory modalities^40,41,42^. Studies show that NE may improve selectivity or increase the magnitude of neuronal responses to sensory stimulation^37^ and promote synaptic transmission and plasticity in threat processing circuits of the amygdala^43^. Indeed, studies have shown that NE signaling through β-ARs in LA and/or BLA can regulate the acquisition^1,2,26^, consolidation^2,29^, reconsolidation^44,45^ and extinction^46,47^ of memory. However, infusion of propranolol into the LA before an expression test does not affect CS-elicited freezing^1^. In contrast, we find a role for NE activity behavioral expression of Pavlovian reactions through CeA. We also note that the null effect of LA propranolol infusion on behavioral expression supports the accuracy of our CeA manipulations.

Studies have shown that optostimulation of LC-NE neurons innervating the BLA can increase anxiety-like behaviors^31^, and sustained (10–15 s) high-frequency (10 Hz) photostimulation caused reversible behavioral arrest^18^. This strong stimulation decreased cortical release of NE, suggesting that brain-wide depletion of NE may be responsible for the observed effects on behavior. It is unlikely that this is the case in our studies with PRS x 8-hM3Dq (Figure 3), which was shown to tonically activate LC neurons at physiological frequencies (~5 Hz)^37^. However, using a manipulation similar to ours, Kane and colleagues show a disruption in a foraging behavior task^48^, possibly due to ‘decision noise’ created by tonic activity of LC that caused behavioral disengagement. This may also be the case in our previous study showing that the invigorating effects of a conditioned stimulus on an avoidance task (Pavlovian-to-instrumental transfer) were disrupted by hM3Dq-mediated excitation of LC^49^. Notably, this negative modulation of active behavioral engagement was also mediated through a LC➔CeA circuit. In terms of pure Pavlovian reactions, however, LC activity and LC➔CeA positively modulate CS-elicited freezing behavior.

Our data suggest that increases in NE itself in the CeA is not sufficient to elicit defensive responses: while the NE LC➔CeA circuit modulated conditioned memory expression, it did not affect unconditioned freezing (Figure 2c, 3c). Although pre-CS freezing was slightly reduced in two experiments (by systemic propranolol (Figure 1b) and systemic injections of CNO in LC hM4Di animals (Figure 2c)), CNO infusion in CeA in hM4Di animals did not change pre-CS freezing levels (Figure 4d). This suggests that the effects of brain-wide blockade of LC-NE, and not LC➔CeA are responsible for this reduction. Notably, baseline freezing was not affected in any of the excitatory hM3Dq manipulations, including the potentiation of β_2_-AR activity with procaterol (Figure 1c), LC activity (Figure 3c) and the LC➔CeA circuit (Figure 5c, 5d). Changes in these would be expected if brain manipulations of NE activity affected unconditioned freezing.

We use chemogenetic manipulations for many of our studies. There has been recent controversy over the use of CNO, which can be metabolically converted into the atypical antipsychotic clozapine^50^. We attempted to circumvent this caveat in several ways. First, we chose systemic doses of CNO that were low enough to minimize this effect^51,52^, and we used mCherry controls that received the same CNO treatments as experimental groups. Furthermore, we supplemented systemic CNO injection studies with direct, intracranial infusion of CNO to bypass the peripheral metabolism of CNO to clozapine.

Studies using other aversive learning paradigms such as inhibitory avoidance also find an important role for NE activity in amygdala, although there are discrepancies when compared to PTC^28-30,47^. For example, NE signaling in the lateral and/or basolateral amygdala is required for memory acquisition in PTC but not for inhibitory avoidance^1,2,28-30,47^. These differences suggest that distinct behavioral and neural processes may underlie each task. Indeed, inhibitory avoidance involves contextual and instrumental learning that are not required for cued PTC^2,28-30,47^. In contrast to lateral and basolateral amygdala, very little is known about NE activity in CeA in both paradigms. One study looking at NE signaling in CeA found no role in the consolidation of inhibitory avoidance memory but did not test for expression effects^29^. Recently we identified a role for NE signaling in CeA in Pavlovian-to-instrumental transfer (PIT), which measures the motivational influence of a conditioned stimulus on active avoidance of a threat^49^. The results suggest that NE activity negatively regulates the invigorating effect of a conditioned stimulus on active instrumental behavior via LC activity and LC projections to CeA^49^. In contrast, in the current study we find a positive modulation of Pavlovian freezing. Taken together, these results suggest that conditioned aversive arousal modulates both Pavlovian and instrumental responses to threat and are consistent with the theory that phasic activation of noradrenergic neurons of the LC, and NE release in CeA, could permit rapid behavioral adaptation to changing environmental imperatives^19,53,54^.

Although LC is a major source of NE to the forebrain^55-57^, other sources of NE to the forebrain include the A1 and A2 adrenergic cell groups in the medulla^58^. Much existing evidence shows A2 adrenergic neurons (located in the dorsal vagal complex, including the nucleus of the solitary tract), also send connections to CeA^58-60^. Our chemogenetic inhibitory studies with hM4Di show subtle but significant effects, which may suggest that other sources of NE, e.g. A2, summate with LC NE in the CeA to positively modulate CS-elicited freezing. Indeed, one study showed that A2 neurons are activated by threat-conditioned stimuli^61^. Future work delineating the relative contribution of these other inputs to defensive responses is warranted.

Given the canonical role of the CeA in expression, in the current studies we focused on this particular phase of conditioning. However, there is some evidence that the CeA itself can mediate Pavlovian memory formation (memory acquisition and/or consolidation)^10,11,62,63^. Recent work exploring the mechanisms underlying this phenomenon suggests that the CeA may convey information about the unconditioned stimulus to the LA during learning^62^. Further studies are needed to determine if NE modulates CeA during the acquisition phase.

Our studies describe the LC➔CeA projection, but a reciprocal CeA➔LC projection has also been reported^64^. This circuit is modulated by corticotrophin releasing factor, which is enhanced by stress and may represent a feedforward excitatory mechanism for the LC-NE-mediated responses to threat. If and how this circuit is involved in defensive reactions is an important subject of future research.

Together with previous studies, the current findings help form a more complete picture of the role of NE in amygdala and emotional learning. Furthermore, amygdala circuits that underlie PTC in rodents and humans are dysregulated in people with fear and anxiety disorders^3,7^. Symptoms of PTSD involve exaggerated responses to stimuli, whether or not it is a threatening^65^, and studies in humans and animals suggest this may due to dysfunction of the amygdala, the LC^66^ and NE activity in amygdala^6,7,22,67-71^. Our current findings support the possibility that a LC➔CeA circuit may underlie exaggerated reactions to stimuli and may explain the efficacy of β-ARs antagonists in some PTSD patients^72-74^.

## Acknowledgments

We thank Cludia Farb, Mian Hou for tissue processing and Dr. Vinod Yaragudri for assistance with imaging, Dr. Hillary Schiff for assistance with behavioral experiments, Dr. Justin Moscarello for discussions during the course of this work, and Dr. Vadim Bolshakov for advice and comments on the manuscript. We thank the University of Pennsylvania Vector Core for packaging the AAV vectors. This study was funded by National Institute of Mental Health (NIMH) grants R01-MH046516 and R01-MH38774, National Institute on Drug Abuse (NIDA) grant DA029053 (J.E.L.).

## Author contributions

Y.G. and R.M.S. designed the experiments, collected and analyzed data, and wrote the manuscript. Y.G., R.M.S and J.E.L. contributed to data interpretation and the final version of the manuscript. All the authors discussed all the results at all stages of the project.

