## Supplemental figures for "A brainstem-central amygdala circuit underlies defensive responses to learned threats"

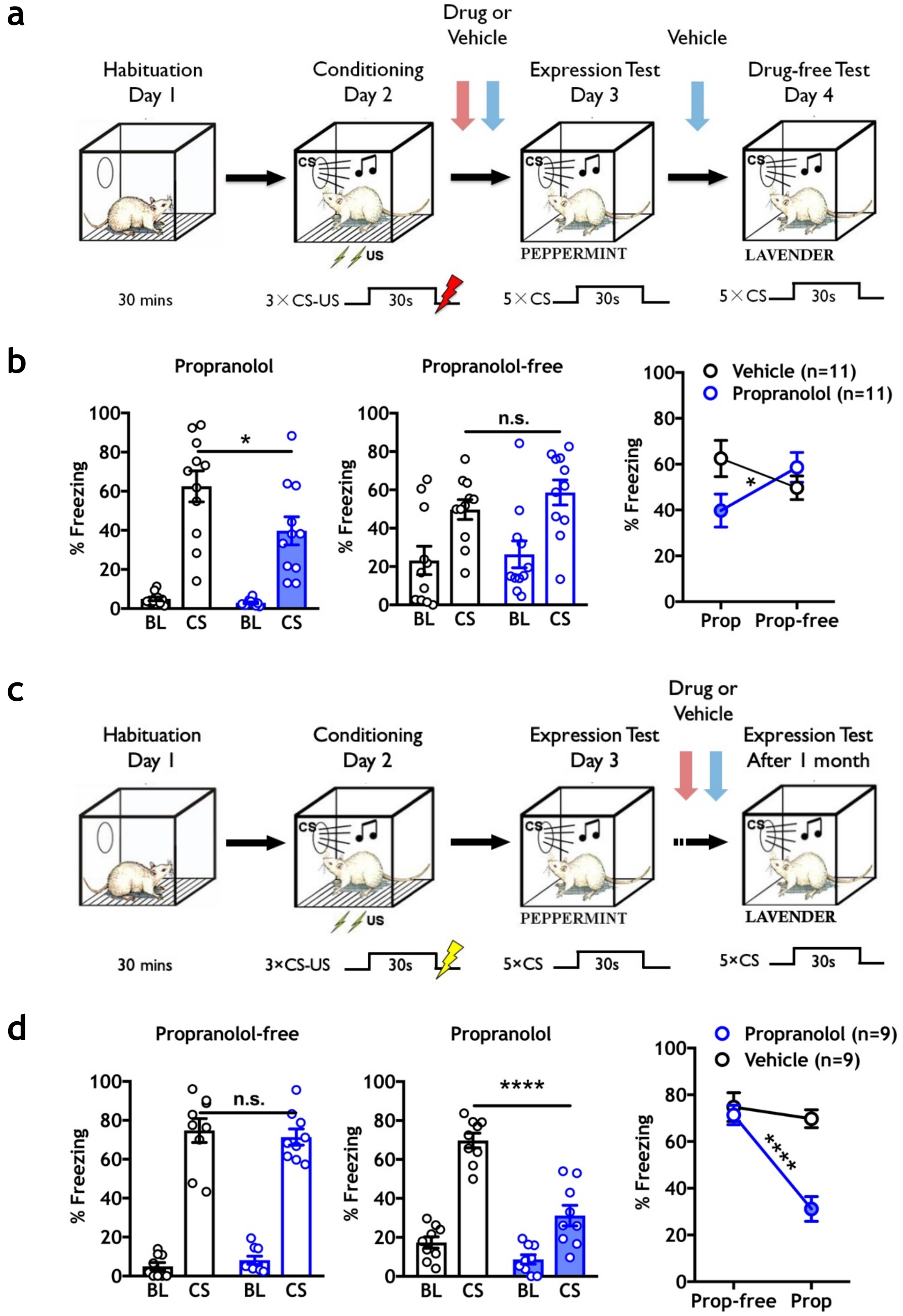

**Supplementary Figure 1. β-ARs is required for Pavlovian defensive responses, regardless of US intensity or memory age. a.** Experimental timeline for strong conditioning paradigm (1.0 mA US). **b.** Propranolol (10 mg/kg) reduced CS-elicited freezing levels during the expression tests compared to vehicle control animals (left panel: n = 11/group, *P = 0.0459), with no effect observed between groups during a drug-free test (center panel, p = 0.2973, n.s.). A within-subject comparison of propranolol versus propranolol-free treatment on CS freezing followed by a post-hoc test revealed a significant difference in CS-elicited freezing for propranolol treatment day versus propranolol-free day (two-way RM ANOVA test, Interaction: F (1, 20) = 9.669, **p = 0.0055; Drug: F (1, 20) = 0.7270, p = 0.4039; Time (Drug vs. Drug-free): F (1,20) = 0.3610, p = 0.5547; Sidak MCS, *p < 0.05 for propranolol treated animals, n.s. for vehicle-treated animals between days). **c.** Timeline of remote Pavlovian threat conditioning (tested one month after conditioning (0.6 mA US)). **d.** Animals showed memory retention after one month, with no difference between groups (left panel, Day 3, n = 9/group; p = 0.6548). Subsequent treatment with propranolol (10 mg/kg, Day 4) significantly attenuated CS-elicited freezing (****p < 0.0001). Within-subject analysis showed a significant difference between a drug-free test (Day 3) freezing and freezing with propranolol treatment (Day 4); two-way RM ANOVA test, Interaction: F (1, 16) = 14.06, **p = 0.0017; Drug: F (1, 16) = 16.55, ***p = 0.0009; Time (Drug-free vs. Drug): F (1,16) = 23.21, ***p = 0.0002; Sidak MCS, ****p < 0.0001 for propranolol treated animals, n.s. for vehicle-treated animals between days. All error bars indicate mean ± SEM. *p< 0.05, **p< 0.01, ***p< 0.001, ****p < 0.0001.

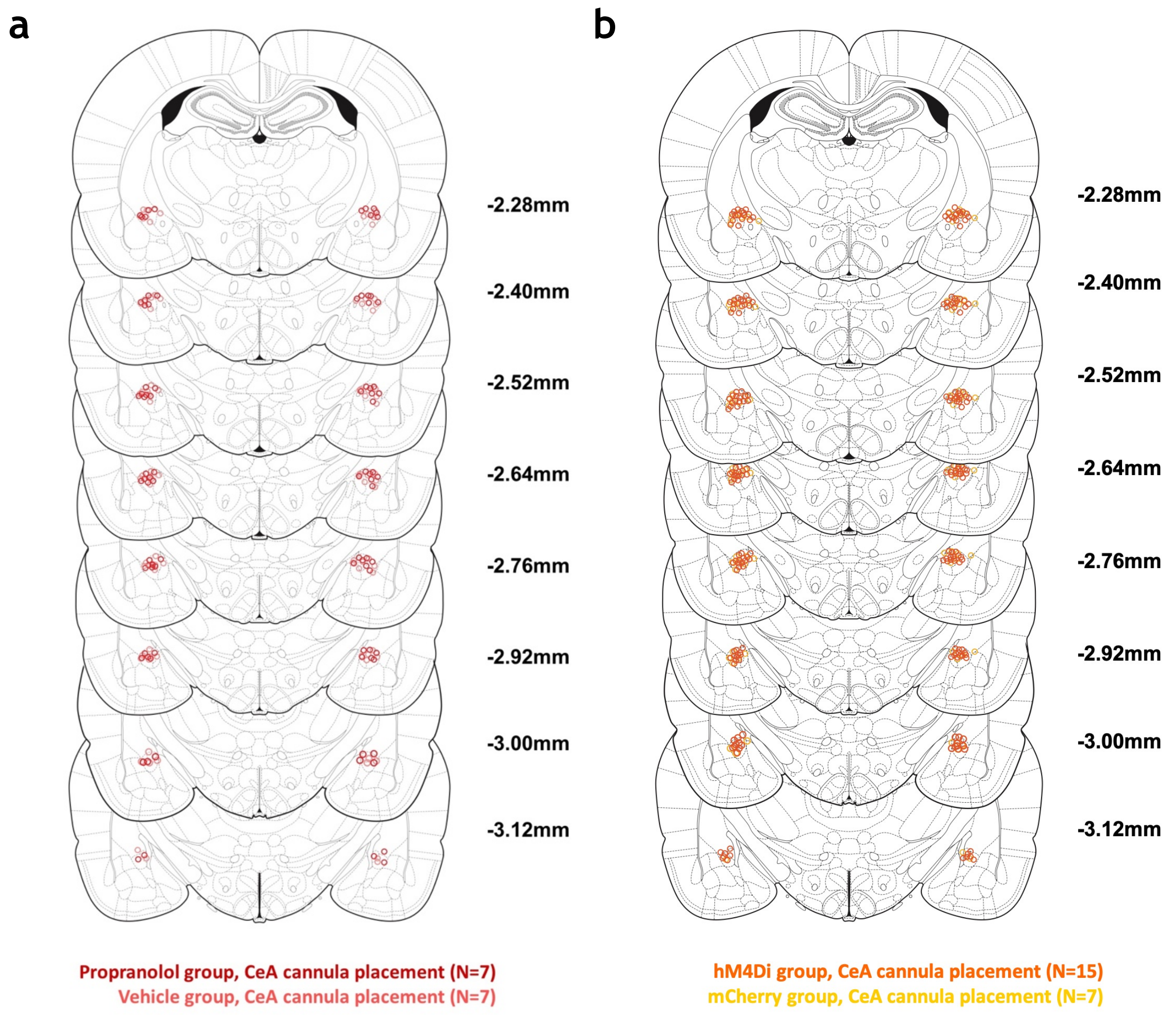
a
b
-2.28mm
-2.40mm
-2.52mm
-2.64mm
-2.76mm
-2.92mm
-3.00mm
-3.12mm
Propranolol group, CeA cannula placement (N=7)
Vehicle group, CeA cannula placement (N=7)
hM4Di group, CeA cannula placement (N=15)
mCherry group, CeA cannula placement (N=7)

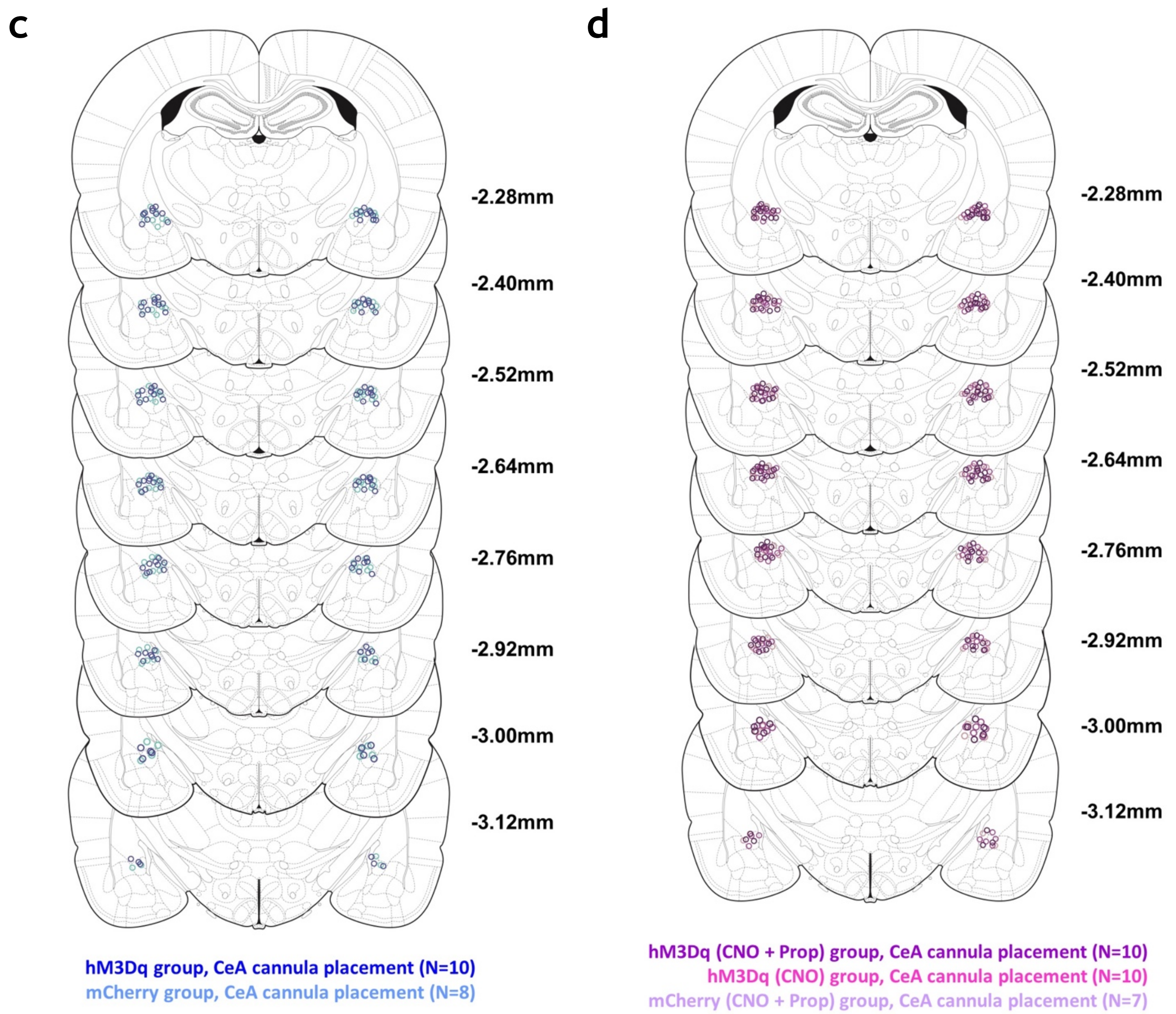


**Supplementary Figure 2. Virus and CeA cannulation targeting for all experiments. Related to Figure 4 and 5.**

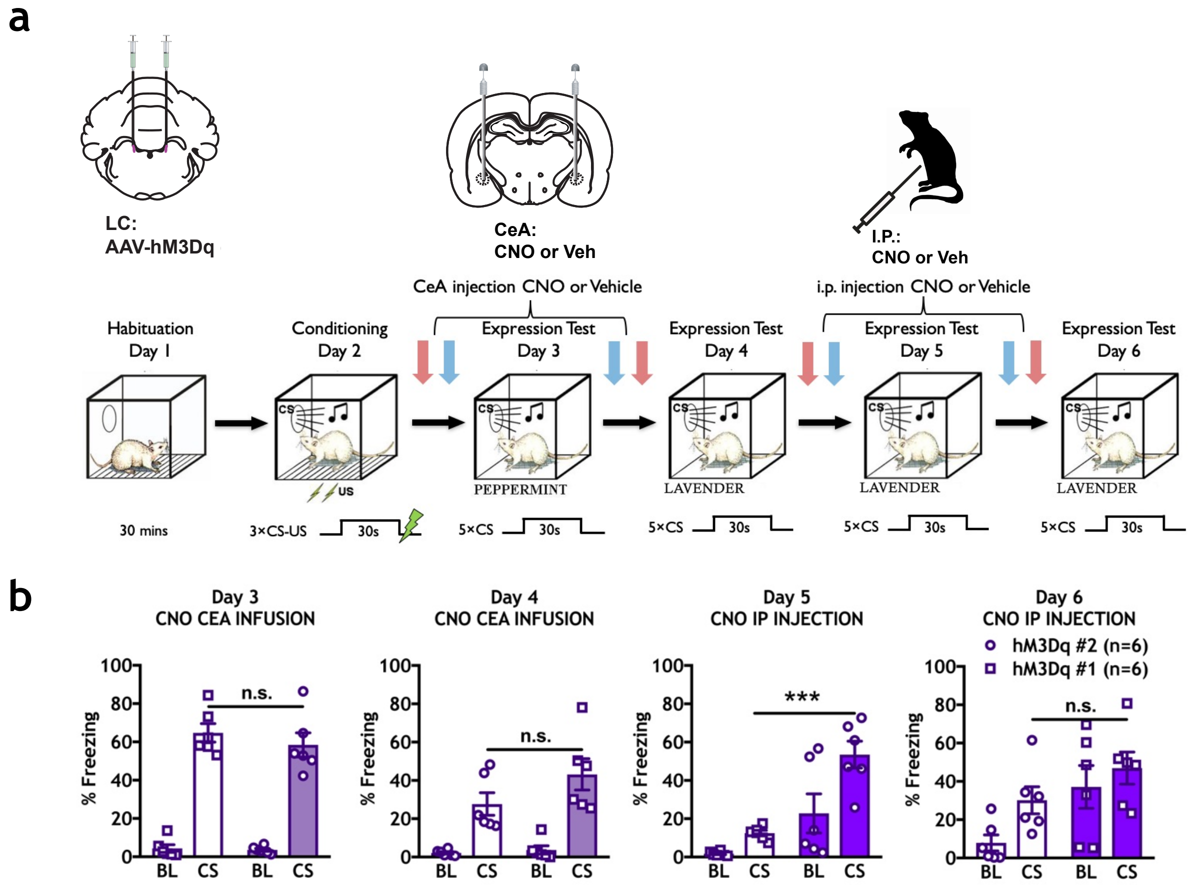


**Supplementary Figure 3. Experiments using axon terminal manipulations require longer incubation periods.**

**a.** Timeline indicating habituation (Day 1), mild conditioning (0.4 mA US) (Day 2), within subject expression test (CeA infusion) (Day 3, Day 4) and within subject expression test (i.p. injection) (Day 5, Day 6) phases. Three weeks following bilateral hM3Dq virus injection in LC, all rats (n=6/group) were trained using a moderate protocol and vertical arrow indicates either CNO (red) or vehicle (blue) treatment strategy. **b.** No significant difference between hM3Dq (CNO) v.s. hM3Dq (vehicle) within CeA infusion on Day 3 and Day 4. Within-subject analysis showed a significant difference in freezing between hM3Dq (CNO) v.s. hM3Dq (vehicle) with systemic treatment (Day 5) (t (10) =5.694, ***p=0.0002). All error bars indicate mean ± SEM. ***p<0.001

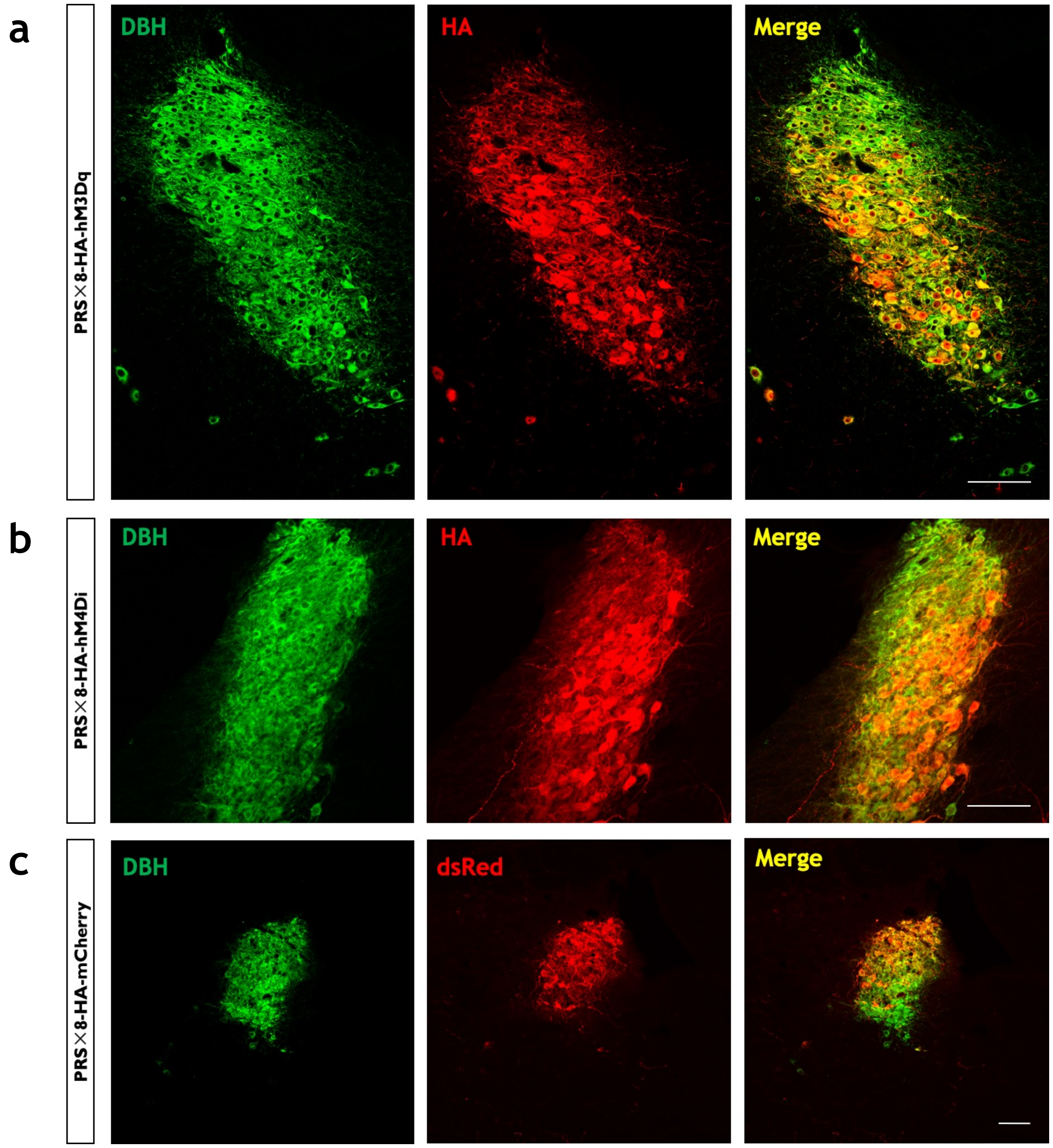


**Supplementary Figure 4. Representative image of DBH$^+$ and HA co-localization in hM3Dq/hM4Di LC rats. a.** Animals injected in LC with AAV-PRS×8-HA-hM3Dq and (**b**) PRS×8-HA-hM4Di show immunoreactivity for HA exclusively in DBH$^+$ neurons. **C.** Animals injected with PRS×8-HA-mCherry (control virus) in LC show mCherry expression only DBH$^+$ neurons. Scale bars, LC panels = 50 µm, respectively.
